# taxMaps - Ultra-comprehensive and highly accurate taxonomic classification of short-read data in reasonable time

**DOI:** 10.1101/134023

**Authors:** André Corvelo, Wayne E. Clarke, Nicolas Robine, Michael C. Zody

## Abstract

High-throughput sequencing is a revolutionary technology for the analysis of metagenomic samples. However, querying large volumes of reads against comprehensive DNA/RNA databases in a sensitive manner can be compute-intensive. Here, we present taxMaps, a highly efficient, sensitive and fully scalable taxonomic classification tool, capable of delivering classification accuracy comparable to that of BLASTn, but at up to 3 orders of magnitude less computational cost. taxMaps is freely available for academic and non-commercial research purposes at https://github.com/nygenome/taxmaps.

## Introduction

Microbial communities of unknown composition can be collected from a wide array of locations. The examination of these microbial communities, known as metagenomics, has become increasingly prominent, with many recent studies focusing on the communities of the human body^1-3^ or from our environment – for example, hospitals^4^, subway stations^5^ or even ATM keypads^6^. High-throughput sequencing enables the unbiased profiling of these communities as well as the ability to investigate clinical samples containing pathogens that are unable to be cultured using traditional laboratory techniques. While the emergence of these technologies has also resulted in more comprehensive databases, querying them in a sensitive manner has become computationally more expensive.

Whether the goal is to estimate the relative abundance or to merely confirm the presence of particular organisms in a given sample, taxonomic classification of each sequence is an essential first step in many metagenomics experiments. Older strategies, either based on machine-learning techniques such as the Naïve Bayes Classifier (NBC)^7^ and PhymmBL^8^ or based on alignment tools such as BLAST^9^, like MEGAN^10^, are slow and do not scale well to the size of today’s experiments. More recently, a new class of faster taxonomic classifiers has emerged. Programs like Kraken^11^ and CLARK^12^ are based on alignment-free strategies where k-mers extracted from the read data are compared to a set of pre-classified k-mers in the database. While these programs can classify millions of reads in just a few minutes, their memory requirements are usually high. Centrifuge^13^ addresses the issue of memory consumption by the use of a FM-index^14^ that allows it to efficiently store and query thousands of bacterial genomes within the memory confines of a standard laptop. This indexing strategy was also employed on the protein homology based classifier Kaiju^15^. While these programs allow for taxonomic classification at an unprecedented speed, no significant improvements in classification accuracy have been reported over Megablast – the least sensitive BLAST program.

Here we describe taxMaps, an ultra-sensitive, customizable and fully scalable taxonomic mapping tool for short-read data designed to deal with large DNA/RNA metagenomics data. taxMaps is designed to facilitate the taxonomic classification operation, featuring thorough preprocessing, the ability to prioritize mapping to multiple indexes, detailed mapping reports and interactive results visualization. Most importantly, by using a novel database compression algorithm that eliminates database redundancy, which improves querying performance and reduces the number of post-querying computations, and an optimal non-exact match mapping strategy, taxMaps delivers classification accuracy that approximates that of BLASTn but in orders of magnitude less time.

## Results

### Database compression

To taxonomically classify short-read data in a comprehensive manner, millions of reads must be compared against DNA/RNA databases, which contain sequences from thousands to millions of organisms. This operation is compute-intensive, given that querying performance is highly dependent on database size and redundancy. This is particularly true when all best hits are to be exhaustively retrieved – something required to ensure maximum classification accuracy. With that in mind, we developed a compression algorithm that eliminates database redundancy by performing a Lowest Common Ancestor (LCA) pre-assignment and collapse for k-mers of length greater than a specified read length (Figure 1a). This allows for non-exact searches to be conducted in the same manner as they would against the original database, resulting in compression that, for the purpose of taxonomic classification, is lossless. Making use of this algorithm we built several databases, some including millions of sequences from more than a million different taxonomic entities (Supplementary Table 1). Compression ratios varied from 1.08 to 4.67, with higher values obtained when using shorter k-mers and usually for databases containing many bacterial genomes, due to the presence of multiple highly homologous strains. Surprisingly, databases compressed at shorter k-mers, despite better compression ratios, require more RAM to be loaded. This may be due to a higher probability of homology between short k-mers, compared to longer ones, which leads to more pronounced sequence fragmentation.

**Figure 1.**
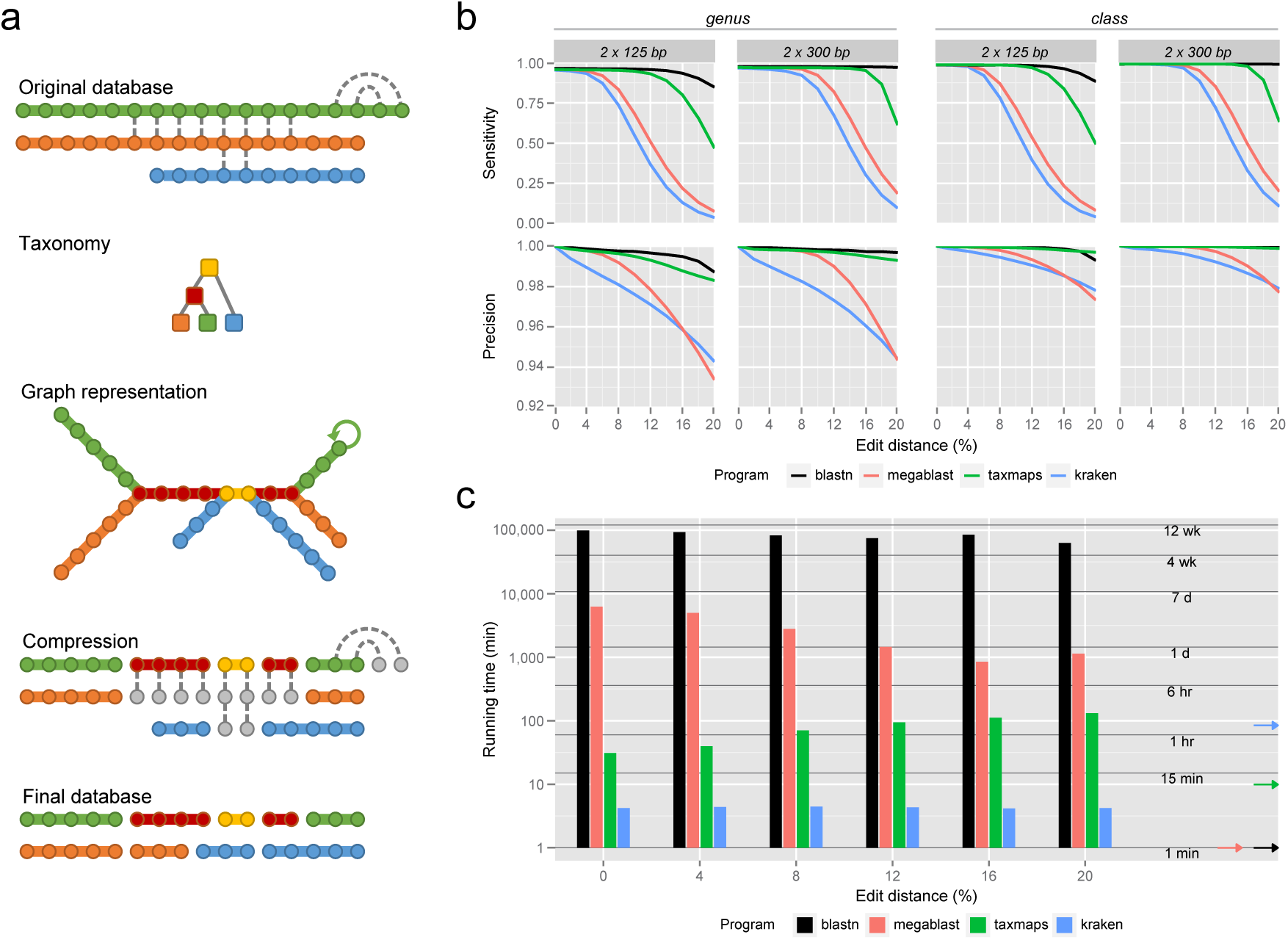
Database compression and classification accuracy and performance on simulated metagenomics paired-end datasets. **(a)** Visual representation of the taxMaps database compression algorithm. Each sequence is represented as an array of k-mers (circles), colored according to their taxon (colored squares). Identical k-mers are linked by a dashed line. During compression, the first instance of every k-mer is reclassified to the Lowest Common Ancestor of all instances of that k-mer in the database while the remaining (grey circles) are disregarded. New sequences, composed of k-mers that share the same taxonomic classification, are assembled on-the-fly as the algorithm transverses the database. A graph representation of the database is also shown. **(b)** Classification sensitivity and precision as function of average sequence divergence and read length at the genus and class ranks. **(c)** Wall clock time required for the classification of six different datasets, each consisting of 10M read-pairs of 125 bp of length, depending on average sequence divergence. The arrows on the right indicate the database loading time for each program.

### Classification accuracy and performance on simulated metagenomes

We compared taxMaps to BLAST (in its two variants Megablast and the more sensitive BLASTn) and Kraken. In this benchmarking exercise, we used NCBI’s nt database^16^ as reference for all four methods to ensure that differences in classification accuracy and speed can only be attributed to algorithmic differences between classifiers and not to reference database differences.

Given that classification accuracy strongly depends on factors such as sequence quality, distance to the closest available sequences in the database, and read length, we have generated 55 simulated paired-end read sets of increasing length (from 75 bp to 300 bp) and divergence (from 0% to 20%) from the reference sequences of more than 4000 different taxonomic units (Supplementary Figure 1).

Classification accuracy results at the genus and class ranks for paired-end reads of length 125bp and 300bp are shown in Figure 1b. It is possible to observe that, while incapable of matching BLASTn accuracy for the most divergent read sets, taxMaps clearly outperforms Megablast and Kraken in both sensitivity and precision. This is particularly striking when sequence divergence is greater than 6%. For instance, on a highly divergent 300bp paired-end dataset (average edit distance = 16%), taxMaps sensitivity and precision at the genus level are 0.951 and 0.995, respectively. On the same dataset, Megablast and Kraken are incapable of classifying more than half of the reads, with sensitivity values of 0.470 and 0.303, at a precision of 0.971 and 0.961, respectively. These results are particularly relevant when choosing the right classifier for metagenomics samples containing organisms that are likely not represented in any database or in situations where the error-rate is high. This trend was observed for all tested read lengths, at virtually all taxonomic ranks for both paired-end and single-end classification (Supplementary Figures 2, 3). Regarding computational performance, Kraken is the fastest method, being capable of classifying 10M 125bp read-pairs in less than 5 minutes, followed by taxMaps that, depending on the average sequence divergence, takes between 31 and 131 minutes to execute the same task (Figure 1c). Nevertheless, this range is comparable to other NGS pipelines (e.g. mapping and variant calling) and 1-2 orders of magnitude faster than Megablast and up to 3 orders faster than BLASTn, on datasets of low sequence divergence to the closest match in the database (Supplementary Figure 4). Interestingly, while for taxMaps the computational cost is positively associated with the average sequence divergence to the reference database, the inverse is true for Megablast, probably because for extreme edit distance values, fewer reads have a seed hit in the database and therefore, no extension operation is performed.

**Figure 2.**
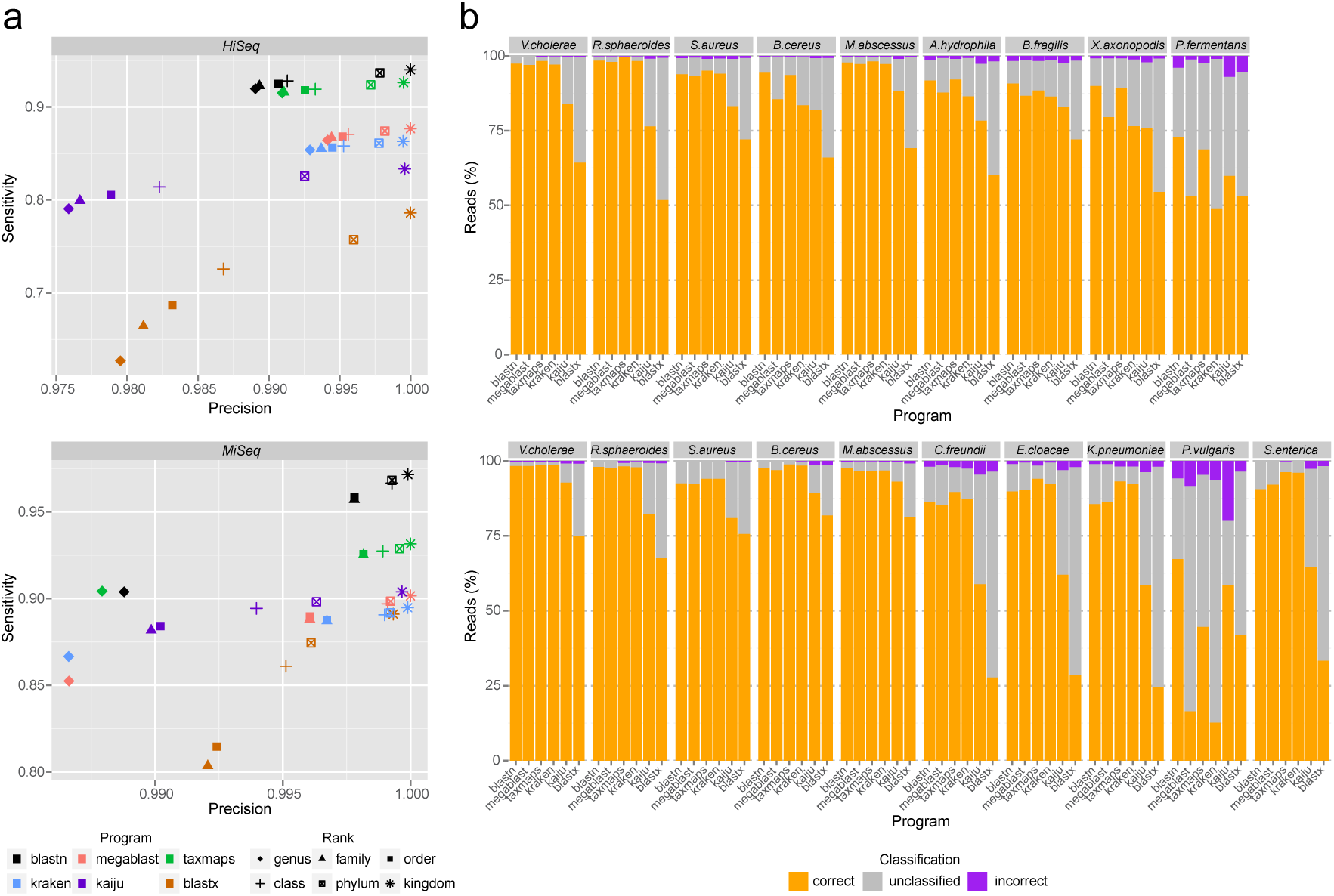
Taxonomic classification accuracy on two mock metagenomics communities. **(a)** Classification sensitivity and precision at six major taxonomic ranks for two real datasets. For visualization purposes, genus-level accuracy values for Kaiju and BLASTx on the *MiSeq* dataset (sensitivity = 0.741 and 0.537, precision = 0.953 and 0.970, respectively) have been omitted. **(b)** The corresponding breakdown per species of the percentage of correct, incorrect and unclassified reads at the genus level.

**Figure 3.**
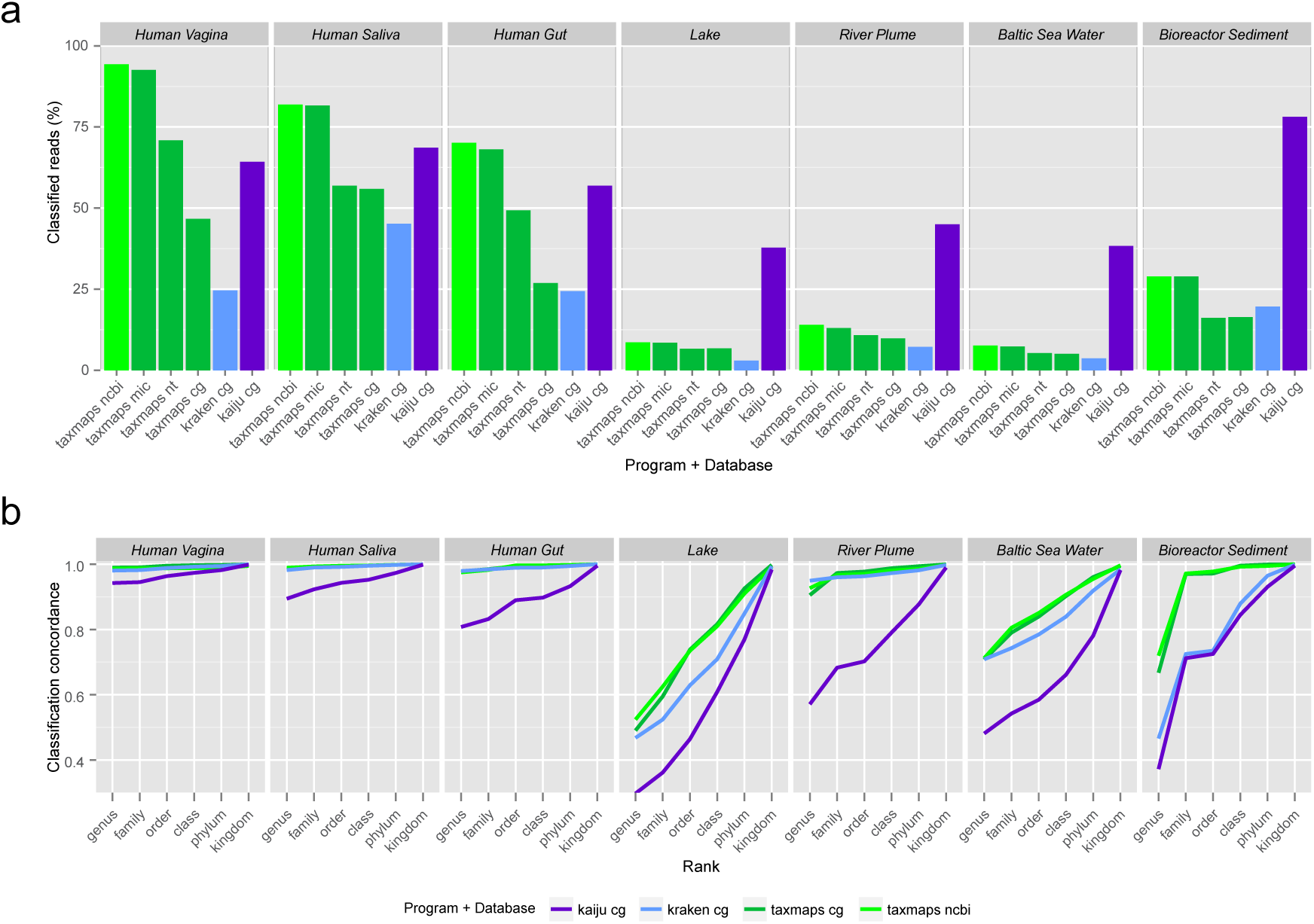
Percentage of classified reads and classification concordance for 7 real metagenomics datasets. **(a)** Percentage of classified reads-pairs for 3 human microbiome and 4 environmental samples. **(b)** Classification concordance between paired mates, as proxy of precision, for 6 major taxonomic ranks.

### Mock communities

To test whether results observed on simulated data would hold when classifying real sequencing data, all four classifiers were tested on two datasets, *Hiseq* and *Miseq*^11^, containing reads from 9 and 10 different bacterial species, respectively. All methods relied on the same database, consisting of complete bacterial, archaea and viral genomes – *refseq_complete_genomes* (see Supplementary Table 1). In this exercise, we also included BLASTx and Kaiju, which rely on protein homology for taxonomic classification. For those, the reference database consisted of all protein sequences annotated on the same set of genomes. For all BLAST methods in this analysis, database search hits were filtered using similar criteria to those used in ^10^. This reduced the number of false-positive classifications originating from small partial alignments, at the cost of some sensitivity (Supplementary Figure 5). As in the results observed for simulated data, BLASTn is the most sensitive method at all the taxonomic ranks considered, with the exception of genus-level classification in the *Miseq* dataset, where taxMaps is marginally more sensitive (Figure 2a). However, it is the least precise of all four nucleotide homology based methods on the *Hiseq* dataset. This discrepancy can be explained by the fact that, on simulated datasets, all reads originate from sequences that are already present in the database, therefore reducing the probability of incorrect classification, while for real sequencing data that is not necessarily the case. Apart from the potential lack of complete genomes in the database, there may be other sequencing artifacts that were not captured in our simulation. After BLASTn, taxMaps is the second most sensitive method at all taxonomic ranks. In fact, for both the *Hiseq* and *Miseq* datasets, taxMaps correctly assigns more reads to the right genus (sensitivity of 0.914 and 0.904, respectively) than any of the remaining programs (Megablast, Kraken, Kaiju and BLASTx) assign to the right kingdom.

For the *Hiseq* dataset, with the default parameter Maximum Edit Distance, *e = 0.2*, taxMaps was slightly less precise at the genus level (0.991) than Kraken (0.993) and Megablast (0.994). It is, however, possible to find values of *e* (*e* <=0.12), where taxMaps is simultaneously more precise and more sensitive than these two methods. On the *Miseq* dataset, running taxMaps with *e = 0.12* drops the genus level sensitivity to that of Kraken but with approximately 60% fewer incorrect classifications (Supplementary Figure 6). Regarding the two protein homology based classifiers, they were the least sensitive and least precise on both datasets at virtually all ranks considered. This result is rather surprising given that protein homology is usually higher than nucleotide homology. One factor that may contribute to this observation is that some reads originate from non-coding regions of the genomes.

While in aggregate BLASTn was the most sensitive method, when further breaking down the results by species (Figure 2b), taxMaps has the highest number of correctly classified reads at the genus level in 5 out of 9 species in the *Hiseq* dataset and in 8 out of 10 species in the *Miseq* dataset (two of which tie with Kraken). For the remaining species, BLASTn obtained the most correct classifications. Interestingly, a few species (*Xanthomonas axonopodis*, *Pelosinus fermentans* and *Proteus vulgaris*), which are divergent from the species in the database, explain most of the differences in overall sensitivity between methods. In those cases, classification performance of taxMaps and BLASTn was significantly higher than that of Kraken and Megablast, being comparable or superior to the protein homology based methods Kaiju and BLASTx, traditionally expected to perform well in that situation.

We also decided to explore, for these two datasets, the taxMaps feature that allows the use and prioritization of multiple databases/indexes. For that, we used the *refseq_complete_genomes* database with a strict value for Maximum Edit Distance (*e = 0.1*) followed by either the *blast_nt*, *refseq_microbial* or *combined_ncbi* databases, with *e = 0.2* (Supplementary Figure 7). While the combination including the *blast_nt* database led to accuracy values similar to those of *refseq_complete_genomes* with *e=0.2*, the use of *refseq_microbial* and *combined_ncbi* raised the genus level sensitivity to values over 0.975 in the *Hiseq* and 0.92 in the *Miseq* datasets, at precision values above 0.991 and 0.989, respectively.

### Human microbiome and environmental samples

While the two mock communities allow for comparisons of classifier accuracy based on real data, they represent a relatively simple classification task, given that most species are well represented in the database used. To assess classifier behavior in a more realistic scenario, we considered 3 human microbiome and 4 environmental metagenomics samples (Supplementary Table 2) as input for Kraken, taxMaps and Kaiju. In this case, due to the large number of reads per sample, we did not consider the slower BLAST methods as they would not represent a practical classification solution. When using the *refseq_complete_genomes* database, Kaiju classified the largest number of reads on all samples, followed by taxMaps and then Kraken, with the sole exception of the Bioreactor Sediment sample, where Kraken classified more reads than taxMaps (Figure 3a). While this suggests that Kaiju may be more sensitive than taxMaps and Kraken on these datasets, the ground truth for these samples is unknown and therefore it is impossible to assess the classification accuracy of each method. To address this problem, we developed a novel rank-level metric called Classification Concordance that, for a given taxonomic rank, can be defined as the percentage of read-pairs where the independent classification of both mates is concordant at that particular rank (see Methods for details). On the simulated dataset described previously, this metric shows a high correlation with classification precision at all ranks individually, and in aggregate (*rho*=0.992) (Supplementary Figure 8). Therefore, it has the potential to be used as proxy for classification accuracy. In Figure 3b it is possible to observe that both taxMaps and Kraken show significantly higher classification concordance than Kaiju on all datasets except for the *Bioreactor Sediment* sample where Kraken classification concordance is only slightly higher than that of Kaiju. This is the sample where, against the general trend, Kraken classified more reads than taxMaps. The lower classification concordance of Kaiju compared to taxMaps and Kraken is particularly striking on the human microbiome samples and the *River Plume* sample, where classification concordance at the phylum level for this classifier is lower than that of taxMaps and Kraken at the genus level. These results suggest that, while Kaiju may classify more reads, it likely does so with much lower precision than taxMaps and Kraken.

Finally, we wanted to investigate how the use of more comprehensive databases in taxMaps would affect the percentage of classified reads and whether there would be a negative effect in classification concordance. We ran taxMaps using *blast_nt*, *refseq_microbial* and *combined_ncbi* databases (Supplementary Table 1) and for all samples the use of these more comprehensive databases resulted in a higher percentage of classified reads. This was particularly clear when using *refseq_microbial* and *combined_ncbi*. Surprisingly, the use of this last database, comprising 374GB of sequence, didn’t have a negative effect on classification concordance compared to *refseq_complete_genomes*, suggesting that taxMaps precision was not affected by the significant increase in the number of sequences in the database. By using very large databases, taxMaps can classify more human microbiome reads than Kaiju and, taking classification concordance as proxy, potentially at much higher precision. As such, taxMaps is particularly appropriate for microbiome studies where maximum classification accuracy at lower taxonomic ranks is desired.

## Conclusions

As genomic databases become more comprehensive, so grows the challenge of how to efficiently utilize such resources to accurately classify the large number of reads generated by high-throughput sequencing technologies. While other recently published methods rely on alignment-free strategies to improve the computational performance of this task, taxMaps’ approach can be considered as an intermediate between that and the more sensitive alignments of BLASTn. By relying on a novel database compression algorithm, taxMaps can conduct very sensitive searches on very large databases while maintaining good performance. Our results using simulated datasets show that the sensitivity and precision of taxMaps approximate that of BLASTn, and are superior to those of Kraken and Megablast, especially as read sequences diverge from the corresponding database reference. These results were further confirmed on the two mock community datasets, where taxMaps delivered the highest number of correct classifications for the majority of the species included. Regarding real metagenomics samples (human microbiome and environmental), when using the same database, both taxMaps and Kraken classified significantly fewer reads than Kaiju. While in a previous benchmark^15^, the number of classified reads has been interpreted as proxy for sensitivity, the ground truth for those datasets is unknown, making it impossible to assess whether classifications are correct or not. To circumvent this limitation, we have developed a novel rank-level metric called Classification Concordance that shows very strong correlation with classification precision. Based on that metric, our results suggest that both Kraken and taxMaps are significantly more precise than Kaiju. Moreover, we show that taxMaps concordance is not affected when using more comprehensive databases that, in the case of the human microbiome samples, led to a significant increase in the number of classified reads.

In summary, our results show that taxMaps offers class-leading accuracy and comprehensiveness while balancing performance, making it uniquely suitable for unbiased contamination detection in large-scale sequencing operations, microbiome studies comprising a large number of samples, and applications where the analysis turnaround time is a critical factor, such as pathogen identification from clinical or environmental samples.

## Methods

### Database creation

Data from the RefSeq Genomes and BLAST nt databases were retrieved through the NCBI FTP server and organized in various databases (see Supplementary Table 1). For each database, duplicate sequence entries were removed and all ambiguous nucleotides converted to N characters. Then, for every distinct k-mer, we computed the LCA between all taxonomic IDs of the sequences containing it, derived from the NCBI Taxonomy database^16^. K-mers were assembled on-the-fly, through extension, into sequences that share the same LCA. This not only eliminates most of the database sequence redundancy, consequently improving mapping performance, but it also significantly reduces the number of post-mapping computations to be performed. This is particularly true for samples containing DNA or RNA from organisms that are highly represented in the databases (e.g. *E. coli*) or for which the repeat content is particularly high. The newly assembled sequences were then indexed (FM-index) using GEM^17^. While the overall strategy is similar to the one employed in Kraken database creation, the fact that this operation is performed on k-mers of length equal or greater than a target read length allows for non-exact searches to be conducted in the same manner as they would against the original database. Thus, this compression is lossless for the purpose of taxonomic classification. The re-assembly of the sequences and use of the FM-index result in a reduction of the memory footprint, allowing for very large databases to be merged and simultaneously queried.

### Classification algorithm

Reads are mapped in single-end mode to an indexed database using GEM mapper^17^, which guarantees that all optimal alignments are retrieved, up to the user defined Maximum Edit Distance (*-e, default = 0.2*) parameter. Each read is then taxonomically classified as the LCA of all database sequences returned. For paired-end classification, reads are classified independently. If the classification of the two ends is discordant, meaning that they are different and the root-to-leaf (RTL) path of one end is not fully included in the RTL path of the other end, the pair is classified as the LCA of both single-end classifications. If the RTL path of one end is contained in the RTL path of the other end, the pair is then classified as the lower taxon of the longest RTL path. In situations where no database match was found for one of the two reads, the pair is classified solely on one read. taxMaps also has a stricter paired-end classification scheme where both ends are required to have database hits. In that scheme, the pair is always classified as the LCA of both single-end classifications, even when one RTL path is contained in the other, ensuring maximum precision at the expense of a higher rank classification.

### Implementation

taxMaps is fully implemented in Python and works as a transparent pipeline-generating script upon user-defined parameters. It reads data in FASTQ format but can also extract unmapped reads from BAM files through Samtools^18^. Processing steps such as adapter removal, low quality end trimming and low complexity filtering are carried out using Cutadapt^19^ and PRINSEQ lite^20^. Users can specify multiple indexes to be queried and define, on an index-specific basis, the maximum edit distance and number of threads used by GEM^17^. Apart from that, taxMaps offers one single-end and two paired-end classification modes (described above). Summary results are given as tables and an interactive report is generated using Krona^21^. Compressed indexes can be downloaded from ftp://ftp.nygenome.org/taxmaps.

### Simulated metagenomics datasets

To build the simulated datasets, we first selected taxa for which the RTL path included all the major taxonomic ranks and had at least one contiguous sequence longer than 100kb in NCBI’s nt database^16^ and then, for each of the 4089 selected taxa (Supplementary Figure 1), we randomly extracted a single 100kb sequence chunk. From these sequences, 55 simulated datasets, each consisting of 10 million read-pairs, were generated using a version of wgsim forked from SAMTools^18^ (https://github.com/lh3/wgsim), by combining five different read lengths (75bp, 125bp, 150bp, 250bp and 300bp) with eleven edit distances (0.0, 0.02, 0.06, 0.08, 0.10, 0.12, 0.14, 0.16, 0.18 and 0.20) and the following additional parameters: fragment length of 550bp, indel fraction of 0.15 and a maximum fraction of ambiguous bases allowed of 0.003. Interleaved FASTQ files were converted to FASTA files for BLASTn and Megablast since these programs were not designed to handle the FASTQ format. Each read ID contains the taxonomic identifier of the sequence from which it was simulated as well as the read length and edit distance of the dataset. All datasets are available at ftp://ftp.nygenome.org/taxmaps.

We ran taxMaps, Kraken, BLASTn, and Megablast on each of the 55 simulated datasets using the NCBI’s nt database as reference for all methods. For BLASTn, the number of read pairs analyzed was reduced to 100,000 by random sampling due to time constraints. Given that BLASTn and Megablast are not taxonomic classifiers per se, the LCA of all best hits for each read was determined. For paired-end classification, the criteria used in taxMaps was applied. To estimate sensitivity and precision, classifications were split into 4 distinct categories: 1) correct, if the correct taxon is included in the RTL path of the assigned taxon; 2) concordant, if the assigned taxon is different from the correct taxon but it is included in the RTL path of the correct taxon; 3) incorrect, if the assigned taxon is not included in the RTL path of the correct taxon nor the correct taxon is included in the RTL path of the assigned taxon; and 4) unclassified, if no taxon was assigned. Rank-level sensitivity is then given by the number of correct classifications at a particular rank over the total number of possible classifications, while rank-level precision corresponds to the number of correct classifications at a particular rank over the number of correct and incorrect classifications at that same rank. Paired-end rank-level sensitivity and precision of each program was calculated at eight major taxonomic ranks (species, genus, family, order, class, phylum, kingdom, and root), for each edit distance and read length combination (Supplementary Figure 2). Similarly, single-end rank-level sensitivity and precision data were also collected for each program from output in single-end mode (Supplementary Figure 3).

In addition to the sensitivity and precision metrics, wall clock time data was collected for each program on all paired-end datasets (Supplementary Figure 4). taxMaps, Megablast and BLASTn were run on a computer cluster running CentOS 7.1 on either Intel Xeon E5-2697 2.60GHz CPUs or Intel Xeon CPU E5-2680 2.80GHz CPUs. Due to the high memory requirements, Kraken was run on a large-memory shared host running CentOS 6.5 on Intel Xeon CPU E7-8830 2.13GHz CPUs. All programs were run using 16 CPUs per job, except for BLASTn, which was run on 8 CPUs given the long-term commitment required of these resources. The wall clock time reported for BLASTn was then extrapolated to match the number of reads classified and numbers of CPUs used by the other programs.

### Mock community datasets

To assess the classification accuracy on real data, we used two mock community single-end datasets, *Hiseq* and *Miseq*, from a previously published benchmark^11^. Each dataset was originally composed of 10,000 single-end reads from 10 different bacterial species. After adapter clipping using Cutadapt^19^, removal of sequences shorter than 31bp and the complete removal of *Streptococcus pneumoniae* from the *Hiseq* dataset due to the presence of chimeric reads that were likely artifacts, there were 8850 and 9953 reads left on the *Hiseq* and *Miseq* datasets, respectively. For each dataset, apart from running taxMaps, Kraken, Megablast and BLASTn, we additionally ran the protein homology based classifiers Kaiju and BLASTx. BLASTx classification followed the same criteria as BLASTn and Megablast. Moreover, a filtering strategy was implemented, for all BLAST programs, using the criteria (minimum bit score, win-score, and top-percent) described by the authors of MEGAN^10^. We selected a win-score of 100 for all programs and minimum bit score cutoffs of 60 for BLASTn and Megablast and 35 for BLASTx. Two values, 5% and 10%, were explored for the top-percent cutoff for BLASTn and Megablast (Supplementary Figure 5). All methods used the *refseq_complete_genomes* database, with the exception of Kaiju and BLASTx that used the correspondent set of annotated proteins.

### Real metagenomics samples

We downloaded 7 Illumina datasets of real metagenomics samples from the Sequence Read Archive (SRA)^22^. Their description and corresponding accession numbers can be found in Supplementary Table 2. On all datasets, adapter sequences were clipped and low quality end bases trimmed (Q<20). Reads were classified with paired-end and single-end modes using taxMaps, Kraken and Kaiju. Due to the high computational requirements, BLAST methods were not considered in this benchmark. For each dataset, apart from determining the number of classified reads by each method, we computed a novel rank-level metric called Classification Concordance. This metric is defined as the percentage of read-pairs for which the independent classification of both ends is either the same or concordant at that particular rank, as long as one of the ends has been classified at that rank or below. For instance, if one end is classified as *Escherichia coli* and the other as Enterobacteriaceae, the classification for that read-pair is considered to be concordant at the species level and at all ranks above. If the second end had been classified as *Proteus vulgaris* instead, the classification would be concordant at the family level and at all ranks above. To assess whether classification concordance could be used as a proxy for precision, we calculated the Spearman’s rank correlation ρ between the two metrics on the simulated datasets, for all methods and at all ranks with the exception of “root”.

## Acknowledgements

This work was partially supported by the Alfred P. Sloan Foundation.

## Authors’ Contributions

AC developed the software. WEC and AC performed the experiments and analysis. AC, WEC, NR and MCZ designed the experiment and wrote the manuscript. All authors read and approved the final manuscript.

